# Timing matters: The temporal representation of experience in subjective mood reports

**DOI:** 10.1101/815944

**Authors:** Hanna Keren, Charles Zheng, David C. Jangraw, Katharine Chang, Aria Vitale, Dylan Nielson, Robb B. Rutledge, Francisco Pereira, Argyris Stringaris

**Affiliations:** Mood Brain and Development Unit, National Institute of Mental Health, National Institutes of Health, Bethesda, MD, USA; Machine Learning Team, National Institute of Mental Health, National Institutes of Health, Bethesda, MD, USA; Emotion and Development Branch, National Institute of Mental Health, National Institutes of Health, Bethesda, MD, USA; Max Planck University College London Centre for Computational Psychiatry and Ageing Research, London, United Kingdom; Wellcome Centre for Human Neuroimaging, University College London, London, United Kingdom

## Abstract

Humans refer to their own mood state regularly in day-to-day as well as in clinical interactions. Theoretical accounts suggest that when reporting on our mood we integrate over the history of our experiences; yet, the temporal structure of this integration remains unexamined. Here we use a computational approach to quantitatively answer this question and show that early events exert a stronger influence on the reported mood compared to recent events. We show that a Primacy model accounts better for mood reports compared to a range of alternative temporal representations, and replicate this result across random, consistent or dynamic structures of reward environments, age groups and both healthy and depressed participants. Moreover, we find evidence for neural encoding of the Primacy, but not the Recency, model in frontal brain regions related to mood regulation. These findings hold implications for the timing of events in experimental or clinical settings and suggest new directions for individualized mood interventions.

**Significance:** How we rate our own mood at any given moment is shaped by our experiences; but are the most recent experiences the most influential, as assumed by current theories? Using several sources of experimental data and mathematical modeling, we show that earlier experiences within a context are more influential than recent events, and replicate this finding across task environments, age groups, and in healthy and depressed participants. Additionally, we present neural evidence supporting this primacy model. Our findings show that delineating a temporal structure is crucial in modeling mood and this has key implications for its measurement and definition in both clinical and everyday settings.

---

Self-reports of momentary mood carry broad implications, yet their underpinnings are poorly understood. We report on our momentary mood to convey to others an impression of our wellbeing in everyday life^1,2^; clinically, self reports of momentary mood form a cornerstone of psychiatric interviewing^3,4^; in research, momentary mood is widely used to quantify human emotional responses, such as in ecological momentary assessment^5-7^. Moreover, theoretical accounts suggest that when we report on our mood, we integrate over the history of our experiences with the environment^8-12^. In this paper we address the fundamental question of the time pattern of this integration—what is the timing of events, e.g. early vs recent—that matter the most for how we report our mood?

The standard account is that momentary mood reporting is predominantly affected by recent reward prediction errors^9^ (RPEs, or how much better or worse outcomes were relative to what was expected). Accordingly, the more surprising an event is (operationalized as a positive or negative RPE) and the more recent it is, the more it will affect momentary mood. In modeling terms, the standard account posits that humans apply a recency-weighting such that their most recent experiences override those in the more distant past (e.g. experiences that happened early in the course of a conversation or a game would matter far less). This computational account has several real-life implications. In terms of measurement of momentary mood, a momentary happiness rating would be a good proxy for one’s most recent experiences. In terms of clinical interactions (such as during an interview or a treatment session) a person’s momentary mood could be lifted by the addition of a positive event.

This standard temporal account is widely applied in models of mood^9-12^, yet is largely unexamined and has not been compared to plausible alternatives. Indeed, at the opposite end of the standard recency model, stands a primacy account of momentary mood. According to a primacy model, experiences that occur early in a conversation, a game or interaction prevail over more recent ones. The intuition for such a model comes from idiomatic expressions such as starting off on the right foot, or empirical evidence which shows that the first instances of an interaction can be highly informative^13,14^. Computationally speaking, early events would be weighted more heavily than recent events, which has several real-world implications. From a measurement perspective, the time scales of momentary mood reporting and of experience would overlap less than in the recency model—the current mood rating would be less of a reflection of the current environment. Moreover, the emphasis on the start of interactions such as interviews or treatment sessions would be much greater.

A computational approach can help us answer this important question as it allows us to make explicit in model terms how humans integrate over their experiences in order to arrive at a self-report of their moods^15^. For this purpose, we developed a novel Primacy model that we pitted against the standard Recency model. We then also examine a host of other plausible models as suggested in disparate literatures about valuation timing^16,17^.

We examine these models across a range of conditions in order to establish their generalizability. First, we examine the different temporal integration of the models in their generalizability across reward environments. To do this we exploited the flexibility of a standard probabilistic task^9^ and adapted it to create different task conditions. a) a *random* environment where there was no consistent trend over time in the direction or value of surprises (RPEs); b) a *structured* environment, where events in the form of RPEs were organized in positive and negative blocks; c) a *structured-adaptive* environment, where the intensity of RPEs was enhanced in real-time to maximize their influence on mood, by compensating for individual and temporal differences in mood response (implemented by adding a closed-loop controller into the standard probabilistic task).

Second, we examine the generalizability of the different temporal integration models across age groups given that previous studies have shown important differences in reward processing particularly between adolescent and adult groups^18-23^.

Third, we also examine the generalizability of the models across healthy volunteers and depressed participants, given the wealth of evidence that depression is associated with aberrations in reward processing^24-27^, which might be also affecting the temporal integration of experiences.

Finally, we compare the neural correlates of key terms of the competing models using whole-brain fMRI. Previous work has shown that the reporting of mood and evoking emotional responses lead to activations in a network of brain areas encompassing the fronto-limbic circuit^9,28,29^. Concordance between computational model parameters and neural activity levels provides evidence that the mechanisms described by the model correspond to the neural processes underlying that behavior. For this reason, we test the correlation of Primacy and Recency model parameters with neural activity measured as BOLD signal during fMRI and then directly contrast between the relations of the two models.

## Results

### The Primacy and Recency models of mood

As a first step we compared the Primacy model versus the Recency model of mood. These were designed to correspond to the general experimental set up that is presented in Fig. 1A and has been used extensively before to answer questions about mood^9,30^. In brief, participants first chose whether to receive a certain amount or to gamble between two values. These values allowed each trial to present to the participant an expectation and an RPE value, where the latter is considered as the difference between the outcome and the expectation values. Subjects were also asked to rate their momentary mood every 2-3 trials, by moving a cursor along a scale between unhappy and happy mood. Such mood ratings have been shown before to correspond to the general state of well-being of participants^30^; we validated this in our dataset with a significant correlation between baseline mood ratings and participant’s depressive symptom scores (t = −3.36, df = 69, p = 0.0012, Cohen’s d effect size = 0.97).

**Fig. 1.**
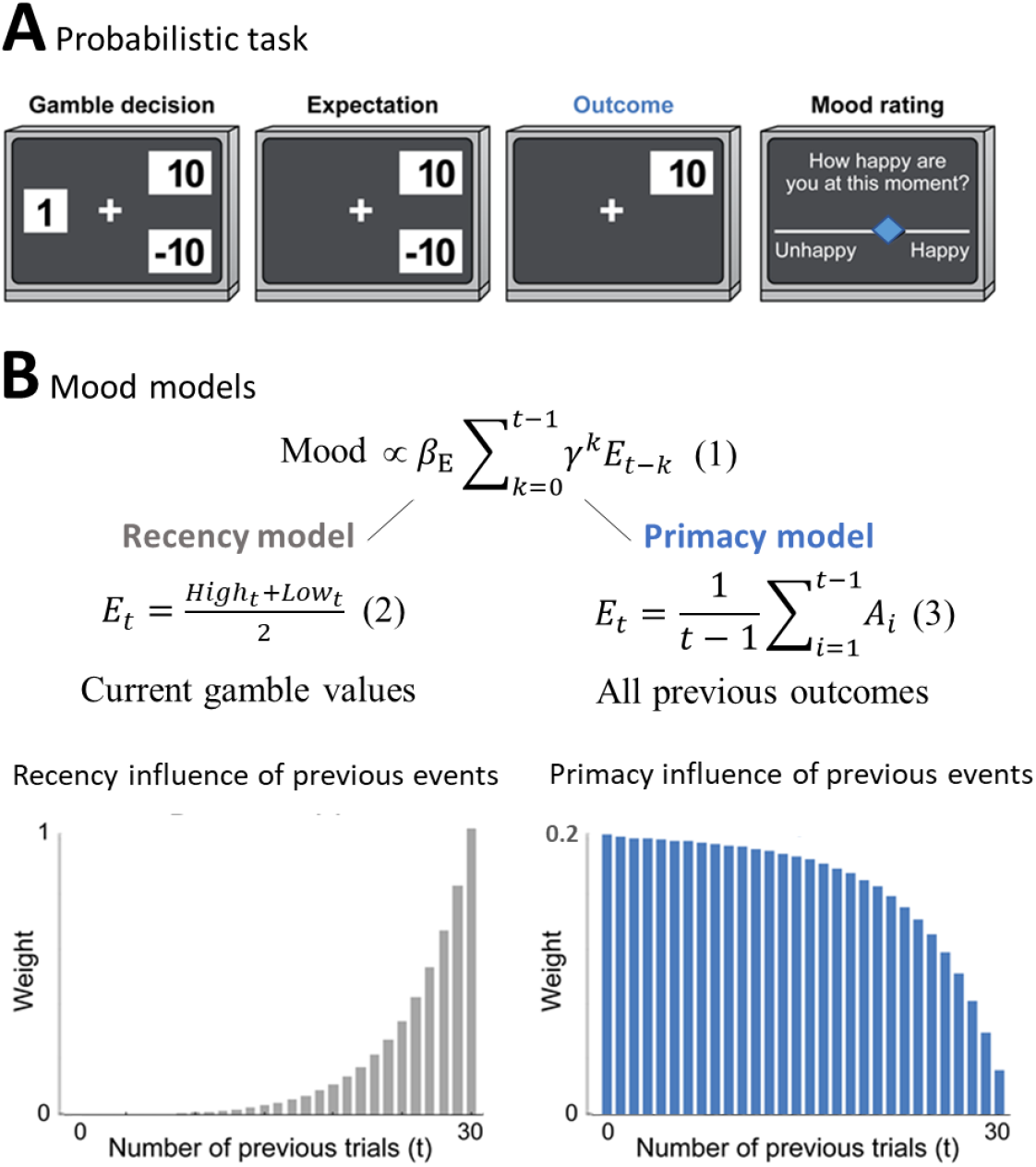
The Primacy versus Recency mood models. (A) Participants played a probabilistic task, where they experienced different RPE values while reporting subjective mood every 2-3 gambling trials. In each trial participant chose whether to gamble between two monetary values or to receive a certain amount (Gamble decision). During Expectation, the chosen option remained on the screen, followed by the presentation of the Outcome value. (B) The expectation term of the mood model is presented in Eq. 1, where *β*_E_ is the influence of expectation values on subjective mood reports. The expectation term of the Recency mood model as developed by Rutledge et al. in 2014^9^ is presented below on the left, where it consists of the trial’s high and low gamble values (Eq. 2). In the alternative Primacy model, as presented on the right (Eq. 3), the expectation term is replaced by the average of all previous outcomes (A_i_). Moreover, as can be seen in Eq. 4-6 in the Methods, the Primacy model has overall fewer parameters compared to the Recency model. The theoretical distributions of the influence of previous events on mood for each model are presented respectively below.

The two principal models, Recency and Primacy, are described in Fig. 1B. Both models consider a cumulative and discounted impact of the expectation term on mood, as shown in Fig. 1B Eq. 1 (Eq. 4-6 in the Methods provide the complete formulation of these models). The Recency Model represents the standard models applied in computational accounts of mood in such setups. In this recency model, expectation is defined as the average between the two gambling values *in the current trial* (Fig. 1 Eq. 2). By contrast, the Primacy Model is our hypothesized account of mood in such setups. In this model, expectation is defined as a *weighted average of all previous outcomes* (Fig. 1 Eq. 3). The critical difference between the two models is illustrated below, by presenting the different theoretical distributions of the influence that events have on mood across the task. As can be appreciated, the Recency model places an emphasis on the most recent trials. By contrast, the Primacy model emphasizes the early ones.

In what follows, we used two main criteria to compare between the models. First, a training error, the Mean Squared Error (MSE) of fitting the model to participant’s mood ratings. Second, a streaming prediction error, a within-subject prediction of each mood rating using the preceding mood ratings. A model performed better if it had significantly smaller error between predicted and rated mood values in these criteria, as tested across participants with a Wilcoxon signed-rank test, with p < 0.05. Moreover, we used a leave-out sample validation and independent confirmatory datasets in all model comparisons.

For completeness, we tested the Primacy model against a range of models with other weighting of past events. Fig. S1 presents a model to which we added to the expectation term a decay parameter and a parameter for how many previous outcomes are included, resulting in various possible distributions of the influence of previous events (Eq. S1-S2).

### Primacy vs Recency model across different reward environments

Here we compare the two models and assess their validity across differently structured reward environments.

#### Random Environment

In order to generate a random reward environment, we used the standard probabilistic task^9^ as described above, where the RPE values were drawn randomly from a pre-defined range of values (Fig. 2A). As shown in Fig. 2B (left panel), this causes mood fluctuations in keeping with previous results (presents the mean across n = 60 participants, with a significant effect of linear time, but not squared time, on mood in a linear mixed-effects model: *β_time_* = −0.31, SE = 0.11, p = 0.006; *β*_*time*^2^_ = 0.0009, SE = 0.002, p > 0.05, and mood change effect size (mean ± SD) = −0.93 ± 1.70).

**Fig. 2.**
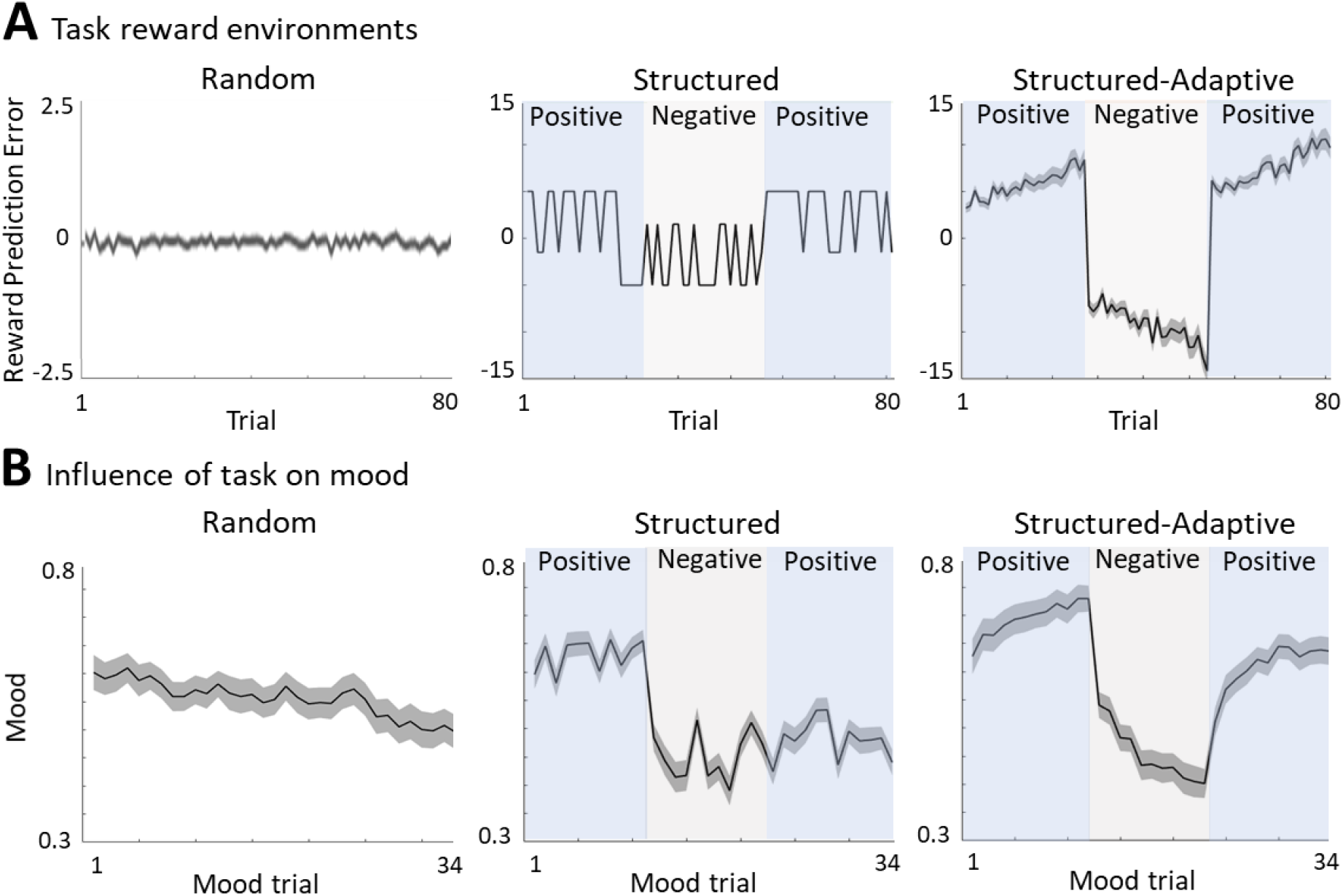
Different experimental reward environments. (A) RPE values received during each task version, averaged across all participants (shaded areas represent SEM). (B) The influence of RPE values on mood reports along the task, averaged across all participants (shaded areas are SEM).

Comparing the two models we found that the Primacy model outperformed the Recency model on the training error criterion (significantly lower MSEs for the Primacy model in a Wilcoxon test with p < 0.05), and performed as well in the streaming-prediction criterion (see Fig. 3 and table S1 for model performance comparison on both criteria).

**Fig. 3.**
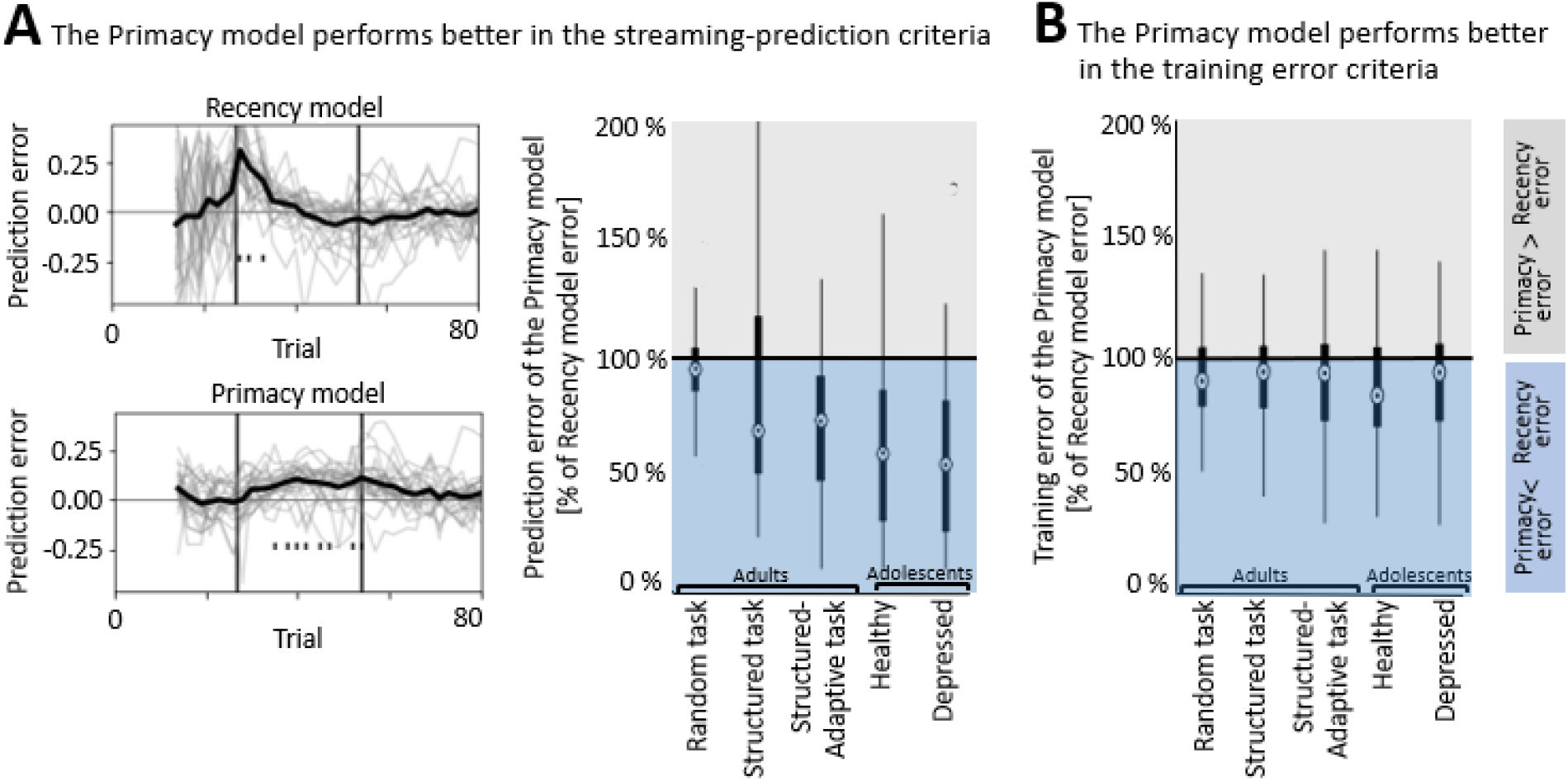
The performance of the Primacy model. (A) Model comparison using streaming prediction criterion, where the model is used to predict each mood rating using the preceding ratings. On the left, the trial level errors in predicting mood with each of the models are shown, for all participants during the Structure-adaptive task (bold line depicts average across all participants), while the right panel presents the median of MSEs of the Primacy relative to the Recency model, in this criterion, across all datasets (edges indicate 25^th^ and 75^th^ percentiles, and error bars show the most extreme data point not considered an outlier). (B) Model comparison using the training error criterion also shows lower MSEs for the Primacy model.

#### Structured Environment

In order to generate an environment that has some structure (consistently positive or negative events), we modified the probabilistic task as shown in Fig. 2A (middle panel): RPE values were divided into three blocks, one of positive RPEs (+ 5), the second of negative RPEs (−5), and a third block of positive RPEs again (+ 5). We found that the experimental set up leads to substantial fluctuations in mood as can be seen in the middle panel of Fig. 2B (presents the mean across n = 89 participants, with a significant effect of time, both linear and squared, on mood in a linear mixed-effects model: *β_time_* = −0.02, SE = 0.006, p < 0.0001; *β*_*time*^2^_ = 0.0004, SE = 0.0001, p = 0.009, and an effect size per block of (mean ± SD): 0.56 ± 1.90, −1.42 ± −1.42, −0.55 ± −0.55, for the first, second and third blocks, respectively).

Comparing the two models we found that the Primacy model outperformed the Recency model on the streaming prediction criteria (significantly lower MSEs for the Primacy model in a Wilcoxon test with p < 0.001), and they had similar training errors (see Fig. 3 and table S1 for model performance comparison on both criteria).

#### Structured Adaptive Environment

Since there can be substantial individual differences in response to events in the environment, we developed a third, adaptive task that tracks individual performance and modifies the environment accordingly. In this paradigm RPE values were not pre-defined but modified in real-time and in an individualized manner, by a proportional-integral (PI) controller^31^, to enhance their potential positive or negative influence on mood over time (see rightmost panel of Fig. 2A for the average RPE values across all n = 80 participants). The task consisted again of three blocks of RPE values, pushing mood towards the highest mood value in the first block, the lowest mood in the second, and the highest mood again in the third block. We found that this experimental setup leads to the largest changes in mood as can be seen in the rightmost panel of Fig. 2B (a significant effect of time, both linear and squared, on mood in a linear mixed-effects model: *β_time_* = −0.04, SE = 0.004, p < 0.0001; *β*_*time*_^2^ = 0.001, SE = 0.0001, p < 0.0001, and an effect size per block of (mean ± SD): 0.92 ± 1.60, −1.75 ± 1.10, 1.45 ± 1.70, for the first, second and third blocks, respectively).

Comparing the two models we found that the Primacy model outperformed the Recency model on both performance criteria, training error and the streaming prediction (significantly lower MSEs for the Primacy model in a Wilcoxon test with p < 0.001; see Fig. 3 and table S1 for model performance comparison).

In what follows, we tested the performance of the Primacy model versus the Recency model, also across different age groups (including different experimental conditions), and across healthy and depressed participants. We then also compared the models by their neural substrates.

### Primacy vs Recency models across different age groups

We found no differences in the strength of the mood changes by age group (no significant group effect on mood in a linear mixed-effects model: F(152,1) = 2.35, p = 0.12, between an adult sample (n = 80) with mean age ± SD = 37.76 ± 11.23, versus an adolescent sample (n = 72) with mean age of 15.49 ± 1.48). The collection of these two data sets differed in age but also in experimental conditions, as the adult sample was collected online (see Methods for details and the pre-registered analysis link), while the adolescent sample was a lab-based collection in an fMRI scanner.

We found that the Primacy model outperformed the Recency model on both performance criteria in both the adult and the adolescent lab-based sample (significantly lower MSEs for the Primacy model in a Wilcoxon test with p < 0.05, see Fig. 3 and table S1).

### Primacy vs Recency models across different diagnostic groups

We found no differences in the strength of the mood changes between the healthy and depressed adolescent participants (when controlling for the difference in baseline mood, there was no significant group effect on mood in a linear mixed-effects model: F(70,1) = 0.77, p = 0.38; between healthy participants (n = 29) with mean ± SD depression score (MFQ) = 1.84 ± 2.49, versus participants diagnosed with Major Depression Disorder (n = 43) with mean depression score of 8.31 ± 6.27; 12 or higher being the cutoff for indicating depression). See Fig. S2 for the distribution of depression scores across this sample.

Comparing the two models in the depressed adolescents sample, we found that the Primacy model outperformed the Recency model on both performance criteria (significantly lower MSEs for the Primacy model in a Wilcoxon test with p = 0.02 for training error and p < 0.0001 for streaming-prediction error; see Fig. 3 and table S1). Moreover, we confirmed this result also in adult participants with high risk for depression (n = 28 participants with CESD scores above 16, being the cutoff for high risk for depression, showed significantly lower MSEs for the Primacy model in a Wilcoxon test with p < 0.001).

The overall better performance of the Primacy model is summarized in Fig. 3. First, we demonstrate the lower trial level errors of the Primacy model in the streaming-prediction criterion, where the consecutive mood rating is predicted based only on preceding mood ratings (shown for all participants during the Structured-Adaptive task in Fig. 3A, left panels). The lower MSEs of the Primacy model in this criterion across all tasks and participant groups, is summarized in the right panel of Fig. 3A. The better performance of the Primacy model in the training error criteria is summarized across all datasets in Fig. 3B. See Fig. S3 for a demonstration of the change over time in task and Primacy and Recency model parameters, shown for a single participant.

### Primacy vs Recency models in relation to brain responses

Finally, we compared the two models on the basis of their relationship to brain activity measured using fMRI. To this end, participants were scanned whilst completing the Structured-Adaptive version of the task. We correlated BOLD signal with the participant-level weights of the Primacy and the Recency model parameters. We found that neural activity preceding the mood rating phase (Fig. 4A), was significantly correlated to the Primacy model expectation term (*β_E_*), which reflects the relationship between mood and previous events (Fig. 4B, cluster at the ACC and vmPFC regions, peak beta = 44.80, t = 3.37, p = 0.0017 corrected to p = 0.05 using 3dClustSim with an ACF function in AFNI as well as an additional Bonferroni correction for comparing three different models). By contrast, the Recency model individual parameters showed no significant relation to neural activity. Next, we directly compared the two models in their relationship to brain activity, by contrasting between the two voxel-wise correlation images (that is, BOLD signal correlation with the Primacy model expectation term weights *β_E_* versus BOLD signal correlation with the Recency model *β_E_*). This showed a significantly stronger relation of the Primacy model to neural activity, specifically in the ACC and vmPFC regions (Fig. 4C, t = 5.00, p = 0.0017).

**Fig. 4.**
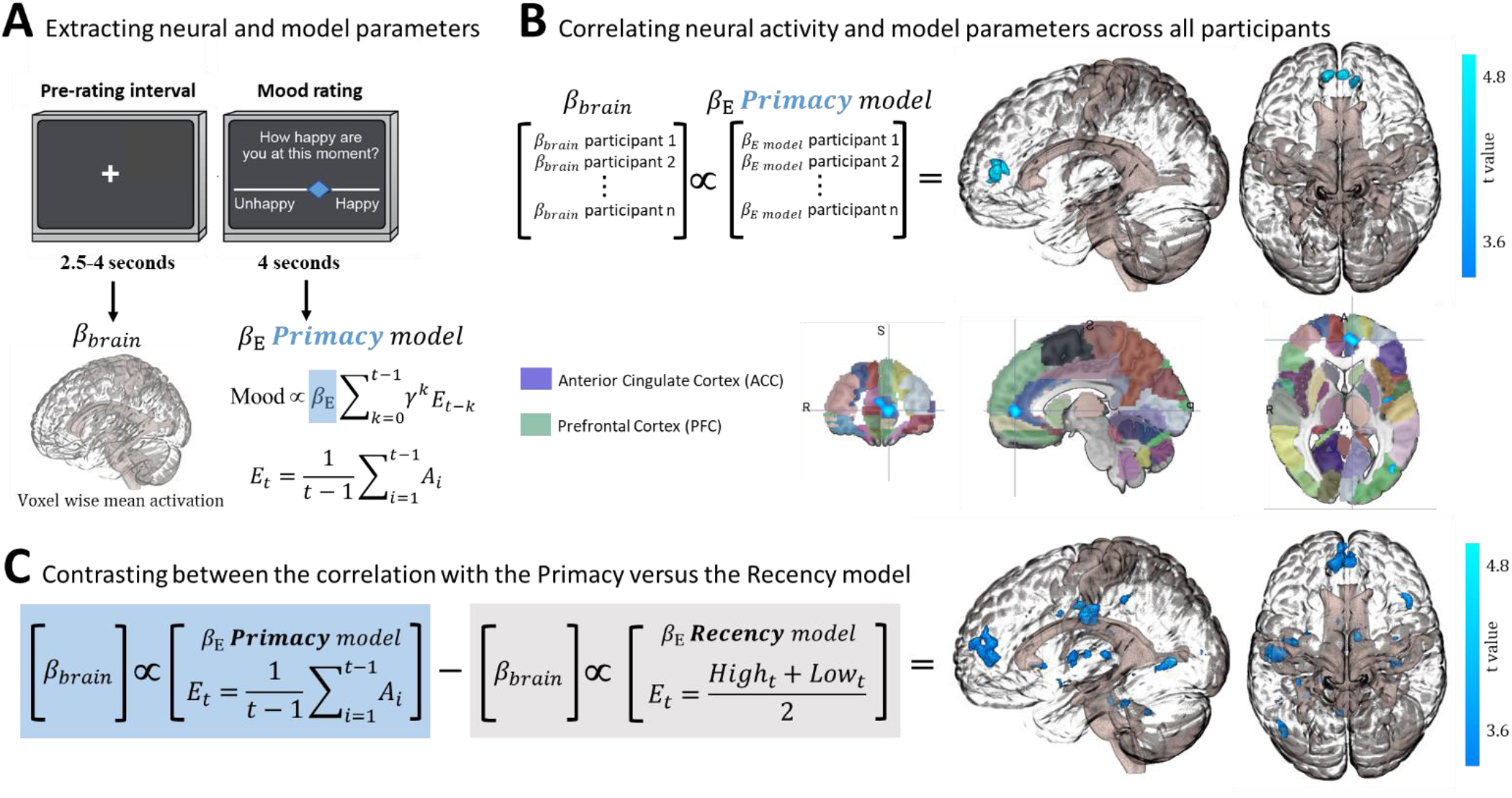
Neural correlates of the Primacy versus Recency model. (A) Extracting individual whole-brain BOLD signal activation maps (*β_brain_*) for the time interval preceding each mood rating (Pre-rating interval), in addition to extracting each individual’s model parameters by fitting mood ratings with the Primacy model (*β_E_*). (B) Correlation across all participants between the individual weights of the expectation term, *β_E_*, and the individual voxel-wise neural activations. A significant cluster was received with a peak at [−3,52,6], size of 132 voxels, thresholded at p ≤ 0.0017 (corrected for multiple comparisons as well as an additional Bonferroni correction for the three 3dMVM models we tested). Below, the resulting cluster of significant correlation presented aligned on the Automated Anatomical Labeling (AAL) brain atlas (focus point of the image is at [−7.17, 50, 4.19] which is located in the ACC region). (C) A contrast between the relation of brain activation with the Primacy versus the Recency model expectation term. This contrast showed a significant difference between the relation of these models to brain signals at [−11,49,9], extending to a cluster of 529 voxels (p < 0.0017).

## Discussion

A fundamental assumption about how humans report on their mood is that they integrate over the history of their experiences. In this paper we sought to test this assumption and establish the temporal structure of this integration.

We show that when humans report on their momentary mood, they do indeed integrate over past events in the environment. However, we find no support for the recency model of integration, i.e. for the standard assumption that most recent events matter the most in this integration. Instead, we find several lines of evidence to support a primacy model of the integration of past experiences in mood reports.

The first line of evidence comes from comparing the primacy, the recency and several other models in a probabilistic task that has been widely used in the past to influence subjective mood reports. This version of the probabilistic task presents participants with random RPE values^9^. The model comparison, conducted using training errors of fitting each model to mood ratings, as well as errors of within-subject prospective prediction of consecutive mood ratings, indicated that a primacy model, i.e. a model that places greater and long-lasting emphasis on early events, described mood ratings the best. The primacy model also outperformed several other plausible models, including a primacy plus recency (U-shaped influence of events) model, an extended model which allowed any timing of previous events to be most influential, as well as models resulting from multiple modifications of both the Primacy and the Recency models (where terms were excluded, merged, or replaced by alternative task values).

We then sought to test whether our findings generalized beyond a random reward environment. We did so in order to emulate real-life situations where negative or positive events tend to cluster over periods of time, such as a conversation between two people that can include consistently pleasant (or unpleasant) events throughout the interaction time frame. We did this by adapting the probabilistic task in two different ways. One, by introducing blocks of predetermined consecutively negative or positive RPEs. In this structured environment, the primacy model outperformed the alternative recency model, in the streaming prediction criterion and performed equally well in the training error criterion. We also modified the probabilistic task in another way, namely by introducing a Proportional-Integral control algorithm. This created a structured adaptive task, with a block of consecutively negative or positive RPEs which were, however, tailored in real-time to each individual’s initial mood level and mood response (such as when we modify our tone of speech during a conversation, according to who we are interacting with and the response we aim for). The primacy model clearly outperformed the alternative recency model and any alternative models, in both performance criteria.

We then also sought to test whether our findings generalized across two important variables. One such variable is age. Substantial evidence shows that adolescence is a time when levels of self-reported mood can change dramatically (e.g. through overall increases in the levels of depression)^18,32-34^. Also, adolescence marks a time when reward processing appears to be different to that of adulthood (with reported increases in the sensitivity of mood)^18-23^. Our primacy model fitted better than the recency alternative in both adolescent and adult samples.

The other variable is subjects’ depression. Its importance is twofold. First, the self-evident fact that persons with depression report a lower mood than non-depressed persons, this is the case in clinical but also in the experimental task setting we and others have used^30^. This difference in mean scores could be reflecting a different way in which persons with depression report on their mood. In keeping with this, the integration of experiences could be happening in a different temporal structure. The second reason is that persons with depression are thought to display reward processing aberrations^24-27^—e.g. in the form of being less sensitive to rewards or learning less from them—that could impact the way in which they integrate across environmental experiences. We address this question specifically in adolescents, the time of a sharp increase in depression incidence^35^, and find no evidence that depressed adolescents applied a different model to the one that their non-depressed counterparts did. Moreover, also adult participants with high scores of depression showed that the Primacy model is a better account of their mood reports. These findings strongly suggest that the temporal representation of experiences offered by the model is robust to important personal characteristics.

We also link the Primacy model to neural activity by a correlation between the model parameters and neural activation at the time preceding mood ratings. Specifically, we show that individual activation at the ACC and vmPFC is correlated to the weight of the expectation parameter of the Primacy model, but not the Recency model. These regions are implicated in mood regulation^28,29,36^’ ^40^ and in underlying decision making relative to previous outcomes^41-44^. Activity in these regions increased as the weight of the expectation parameter *(β_E_*) of an individual was higher. Since the weight of this parameter determines the influence of previous outcomes on mood, this result suggests that these regions’ activity plays a role in mediating the integration of previous outcomes to a subjective mood report. Therefore, the strength of this model-based fMRI analysis^45,46^ is in allowing us to link neural signals to the computational relation between previous experiences and subjective mood reports.

Our experiments examine only a short space of time, no longer than 40 minutes. Human experiences are undoubtedly integrated over longer time periods, including temporally distant events in childhood. Whilst these are inherently difficult to model experimentally, it is noteworthy that early life experiences, such as early adversity, are thought to exert long-term influences on mood^47-49^. We also note that the time scales of our experiments are congruous to a number of real-life situations, both in research and clinical terms.

In research terms, self-reported mood in ecological momentary assessment (EMA) is typically within the span of hours^5-7^. Given that the goal of EMA is often to uncover mood dynamics in relation to experiences in the environment, our results strongly indicate that explicit modeling of the relative timing of these two variables to each other may be crucial. Similarly, during fMRI and other scanning, researchers often ask participants to report on their mood during these sessions and use these to relate to neuroimaging results. Our results suggest that not just the value of events as such (whether, for example, an aversive film was shown to participants), but also when it was shown may differentially impact such reports.

In terms of clinical events—such as patients’ interactions with healthcare professionals for the purposes of psychotherapy or medication treatment—these typically last for about an hour. Importantly, the assessment of treatment progress relies on self-report (or clinician assessment of patients’ reports). Our results suggest that timing of such reports in relation to experiences during treatment could be an important source of variance.

Moreover, although our experiments test for different temporal structures of reward, they use a single type of task, a simple gambling decision task. It might be that the temporal structure of mood dependence is sensitive to the type of task or the context (i.e. social situations such as a conversation, may include a different integration), which is an important matter for future studies. Another potential caveat relates to the online data collection using the Amazon Mechanical Turk (Mturk) platform, which can possibly include a different subset of participant characteristics^50^, but importantly, our results were robust to this difference and were well-replicated in our lab-based participants. In respect to the Primacy model characteristics, we aimed to minimize the divergence from the existing Recency model (we therefore changed the expectation term to consider the average of all previous outcomes but maintained the sum and the overall exponential discounting of that term). This computational modification between the models reflected our hypothesis that expectations in non-random temporal structures of rewards would be also influenced by the history of previous outcomes. This modification then resulted in a Primacy weighting. Nevertheless, the better performance of a Primacy weighting was consistent also when considering other formulations for weighing of previous events (and without taking into account the fewer parameters of the Primacy model).

More generally, our findings point to the importance of studying the temporal architecture of the interplay between experiences and mood. So far, computational accounts of mood have focused on event value (either in terms of expectations or outcomes or both) as influences on subjective reports of wellbeing, while neglecting the importance of time^51,52^. Our results suggest that in addition to these influential properties of the environment, the dimension of time, i.e. the temporal structure of previous events, also plays an important role, and that rather than being a matter of what happened most recently, the temporal representation of experience in mood seems to be dominated by a long-lasting effect of early events.

## Methods

### 1. The Primacy versus Recency mood models

The formulation of both models consisted of two dynamic terms: the expectation term (E) and the RPE term (R), which is the difference of the outcome relative to the expected value.

Specifically, the Recency model of mood at trial t (M_t_) was defined as:

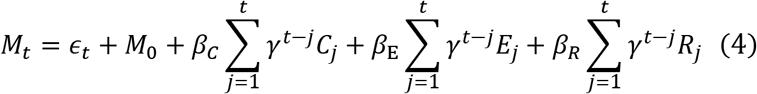

where *ϵ_t_* is a random noise variate with some unknown distribution (we may assume it to be Normal with mean 0 and standard deviation *σ*), *M*_0_ is the participant’s baseline mood, *γ* ∈ (0,1) is an exponential discounting factor, C_j_ is the non-gamble certain amount at trial j (if not chosen then C_j_ = 0 and when chosen instead of a gamble then E_j_ = R_j_ = 0), *β_c_* is the participant’s sensitivity to certain rewards during non-gambling trials, *β_E_* is the participant’s sensitivity to expectation and *β_R_* is the sensitivity to surprise during gambles.

In this model the expectation term at trial t (E_t_) was defined as the average between the two gamble values (see Fig. 1 Eq. 2) and the RPE term, R, was defined as:

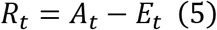

A_t_ being the trial outcome.

In the Primacy model, the expectation term was replaced by the average of all previous outcomes (Fig. 1 Eq. 3) and R was defined similarly as shown in Eq. 5 above. The overall Primacy model for mood at trial t was:

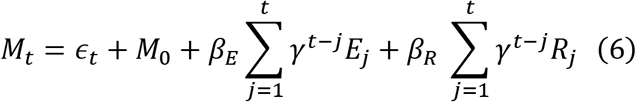

where *β_E_* and *β_R_* are the participant’s sensitivity to previous outcomes and to how surprising these outcomes are relative to expectation, respectively. Note that this model performed better when we did not distinguish between gambling and non-gambling trials, which was another divergence from the standard Recency model.

### 2. Model fitting

All models were fit using a TensorFlow package. We chose group regularization constants by creating simulated datasets with realistic parameters and selecting the regularization parameters from a grid that had the best performance. The grid consisted of powers of 10 from 0.001 to 10000. For optimization, we used the following generic parametric model across subjects:

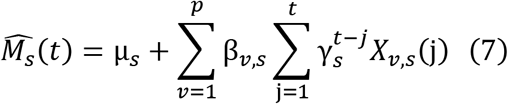

where s indexes the subject, M_s_(t) is subject’s mood rating at trial t, μ_s_ is the subject-specific baseline mood, v is one of p time-varying task variables X (e.g., expectation or RPE values at each trial j), and β_v,s_ are subject-specific coefficients for each time-varying variable X_v,s_ (note that we constrain β_1_,…,β_3_ ≥ 0).

To facilitate optimization, we further re-parameterized the discount factor γ_s_ by defining

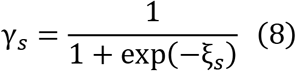

so that *ξ_s_* is an unbounded real number.

We found that the use of group-level regularization was necessary in order to stabilize the estimated coefficients. This took the form of imposing a variance penalty on *ξ* and on each coefficient βv. The empirical variance is defined as

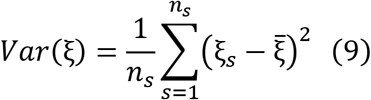

where n_s_ is the number of subjects, and 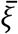 is the group mean:

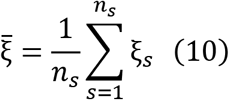

We define Var(X_v_) for v = 1,…,p likewise.

The objective function is therefore:

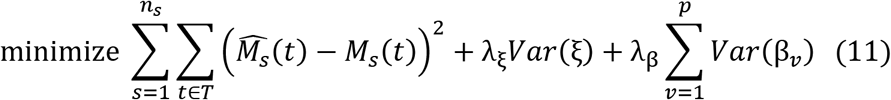

where T is the set of trials where M_s_(t) was defined (optionally, one can also discard the first few trials in T to minimize window effects, we required t ≥ 11), with λ_ξ_ = 10 and λ_β_ = 100 as the regularization parameters with the best performance in recovering the simulation ground truth for both models.

A leave-out sample validation approach was used in all model fitting, where a sub-sample of 40% randomly selected participants were modelled and then results were confirmed on the entire sample.

### 3. Model comparison

The criteria for comparison between models were:

1. The model fit MSE, the error of fitting the model to participant’s mood ratings.
2. Streaming prediction MSE, the error of predicting the t-th mood rating using the first t-1 mood ratings.

In both criteria, lower MSE were tested for significance using the one-sided Wilcoxon signed-rank test. We chose a one-sided null because the conservative null would be that the new approach is equal or worse than the existing approach. The Wilcoxon signed-rank test tests the null hypothesis that two related paired samples come from the same distribution.

### 4. Testing the Primacy model across reward environments

#### The Random task

Participants played a gambling task where they experienced a series of different RPE values while rating their mood after every 2-3 trials. Each gambling trial consisted of three phases: (1) Gamble choice: 3 seconds during which the participants pressed left to get the certain value or right to gamble between two values (using a four-button response device); (2) Expectation: only the chosen certain value or the two gamble options remained on the screen for 4 seconds. (3) Outcome: A feedback of the outcome value was presented for 1 second, followed by an inter-trial-interval of 2-8 seconds. The certain value was the mean of the two gamble values. Participants completed 81 trials.

The mood rating consisted of two separate phases: (1) Pre-rating mood phase, where the mood question: “How happy are you at this moment?” was presented for a random duration between 2.5-4 seconds, while the option to rate mood was still disabled; (2) Mood rating by moving a cursor along a scale labeled “unhappy” on the left end and “happy” on the right end. Each rating started from the center of the scale, and participants had a time window of 4 seconds to rate their mood. The cursor could move smoothly by holding down a single button press towards the left or the right directions. Each rating was followed by a 2-8 seconds jittered interval. Participants completed 34 mood ratings and the task lasted 15 minutes.

#### The Structured task

In this version participants experienced blocks of the same high or low RPE values, i.e., stable patterns of positive or negative events. The task consisted of three blocks, where each block had 27 trials and 11-12 mood ratings. Trials were identical to the Random version in appearance and timing but differed in having pre-defined gamble and outcome values, in order to generate blocks of pre-defined RPE values. Participants again completed overall 34 mood ratings in 15 min of task time.

#### The Structured-Adaptive task

The Structured-Adaptive version was designed to maximally influence mood upwards or downwards, by increasing or decreasing RPE values in real-time. This task was identical to the Structured task in the block design and number of trials but differed in RPE values being calculated in real-time using a closed-loop control (Proportional-Integral algorithm used in control of non-linear systems in engineering^31^). Specifically, following each mood rating, RPE values were increased or decreased according to the difference of mood from the target mood, which was set to the highest mood value in the first and third positive blocks and to the lowest mood during the second block. This setup therefore generated personalized “reward environments”, as the task values were calculated online according to individual mood response and were not pre-determined as in conventional paradigms.

#### Participants

Participants completed either the Random task (n = 60, mean age ± SD = 39.81 ± 13, 44% females), the Structured task (n = 89, mean age ± SD = 37.55 ± 10.46, 44% females), or the Structured-Adaptive task (n = 80, mean age ± SD = 37.76 ± 11.23, 42% females). These participants were recruited from Amazon Mechanical Turk (MTurk) system and completed the tasks online. Analyses of this Structured-Adaptive dataset were pre-registered on OSF, to confirm our modelling results (https://osf.io/g3u6n/). The MTurk Worker ID was used to distribute a compensation of $8 for completing the task and a separate task bonus between $1 to $6 according to the points gained during the task. Participants were instructed before the task that they would receive a payment which is proportional to the points that they gain during the task. These study populations were ordinary, non-selected adults of 18 year of age or older. Participants were not screened for eligibility, all individuals living in the US and who wanted to participate were able to do so. Participants were restricted to doing the task just once. Three participants were excluded from analyses due to an error in the task script where mood ratings were inconsistently spread along the 3 blocks. All participants received similar scripted instructions and provided informed consent to a protocol approved by the NIH Institutional Review Board.

#### Statistical testing of the influence of reward environments on mood

We applied a linear mixed effects model to estimate the task influence on mood, using the nlme package in RStudio (2020). This model enabled the estimation of the across-participants significance of mood change while controlling for the within-participant variability in mood change slopes and intercepts, defined as random effects. Specifically, the independent variable was the response variable of interest mood (M), and the dependent variables were time (t, which is the trial index) and time squared (t^2^), with the two different time variables considered as random effects), as follows:

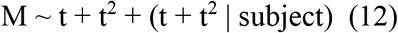

The effect was considered significant with p < 0.05. All t-tests conducted were 2-sided.

### 5. Testing the Primacy model across participant characteristics and in relation to neural signals

**A**n additional dataset of the Structured-Adaptive task was collected in an fMRI scanner, providing us with different experimental conditions, a different age group of adolescent participants, data of participants diagnosed with depression, and a recording of neural signals during the task (n = 72, mean age ± SD = 15.49 ± 1.48, 76% females, mean depression score MFQ ± SD = 5.81 ± 5.98, n = 43 participants met diagnostic criteria for depression according to DSM-5, of whom at the time of the experiment n = 18 had an ongoing depressive episode and n = 35 were medicated). These participants completed the task in an fMRI scanner and were compensated for doing the task and for scanning, as well as receiving a separate bonus proportional to the points earned during the task (a value between $5 to $35). This task version lasted 24 min instead of the duration of 15 min of the online versions, to allow for an optimal analysis of brain data. Participants were screened for eligibility and inclusion criteria were the capability to be scanned in the MRI scanner and not satisfying diagnosis criteria for disorders other than depression according to DSM-5. Overall, five participants were excluded from analyses due to incomplete data files, and three additional participants were excluded due to repeatedly rating a single fixed mood value for an entire block of the task. These participants received the same scripted instructions and provided informed consent to a protocol approved by the NIH Institutional Review Board.

### 6. Analyzing the neural correlates of the Primacy model

#### fMRI data acquisition

Participants in the adolescent sample, performed the Structured-Adaptive task while scanning in a General Electric (Waukesha, WI, USA) Signa 3-Tesla MR-750s magnet, being randomly assigned to one of two similar scanners. Task stimuli were displayed via back-projection from a head-coil mounted mirror. Foam padding was used to constrain head movement. Behavioral choice responses were recorded using a hand-held Fiber Optic Response Pad (FORP). Forty-seven oblique axial slices (3.0 mm thickness) per volume were obtained using a T2-weighted echo-planar sequence (echo time, 30 ms; flip angle, 75°; 64 × 64 matrix; field of view, 240 mm; in-plane resolution, 2.5 mm × 2.5 mm; repetition time was 2000 msec). To improve the localization of activations, a high-resolution structural image was also collected from each participant during the same scanning session using a T1-weighted standardized magnetization prepared spoiled gradient recalled echo sequence with the following parameters: 176 1-mm axial slices; repetition time, 8100 msec; echo time, 32 msec; flip angle, 7°; 256 × 256 matrix; field of view, 256 mm; in-plane resolution, 0.86 mm × 0.86 mm; NEX, 1; bandwidth, 25 kHz. During this structural scanning session all participants watched a short neutral-mood documentary movie about bird migration.

#### Data preprocessing

Analysis of fMRI data was performed using Analysis of Functional and Neural Images (AFNI) software version 19.3.14^48^. Standard pre-processing of EPI data included slice-time correction, motion correction, spatial smoothing with a 6-mm full width half-maximum Gaussian smoothing kernel, normalization into Talairach space and a 3D non-linear registration. Each participant’s data were transformed to a percent signal change using the voxel-wise time series mean blood oxygen level dependent (BOLD) activity. Time series were analyzed using multiple regression^49^, where the entire trial was modeled using a gamma-variate basis function. The model included the following task phases: Gamble choice: an interval that lasted up to 3 seconds, from the presentation of the 3 monetary values to the choice button press, left for the certain amount or right to gamble. Expectation: a 4 seconds interval from making the choice of whether to gamble to receiving the gamble outcome. Outcome: a 1 second interval during which the received outcome is shown. The Pre-rating interval: a variable interval between 2.5-4 seconds, when the mood question is presented but the option to rate mood is still disabled. Mood rating phase: a 4 seconds interval during which participants rate their mood. The model also included six nuisance variables modeling the effects of residual translational (motion in the x, y and z planes), rotational motion (roll, pitch and yaw) and a regressor for baseline plus slow drift effect, modeled with polynomials (baseline being defined as the non-modeled phases of the task). Echo-planar images (EPI) were visually inspected to confirm image quality and minimal movement.

Statistical significance was determined at the group level using 3dClustSim (the latest acceptable version in AFNI with an ACF model) which generated a corrected to p < 0.05 voxel-wise significance threshold of p < 0.005 and a minimal cluster size of 100 voxels. We analyzed relation to model parameters with neural activity during three different phases of the task: activation during the pre-mood rating period, mood rating encoding (with mood values as a parametric regressor of the mood pre-rating period), and task based RPE encoding (RPE values as a parametric regressor of the outcome period). Since these are three separate tests, we added a Bonferroni correction to the multiple comparison correction, which resulted in a final p-value threshold of 0.005 / 3 = 0.0017. We then ran a whole-brain, group-level ANOVA (3dMVM^53^ in AFNI) with the weights of the Primacy or the Recency model as between-participant covariates of each of these neural activations (each participant’s neural activity was represented by a single whole-brain image of activation across all trials).

---

**Fig. S1.**
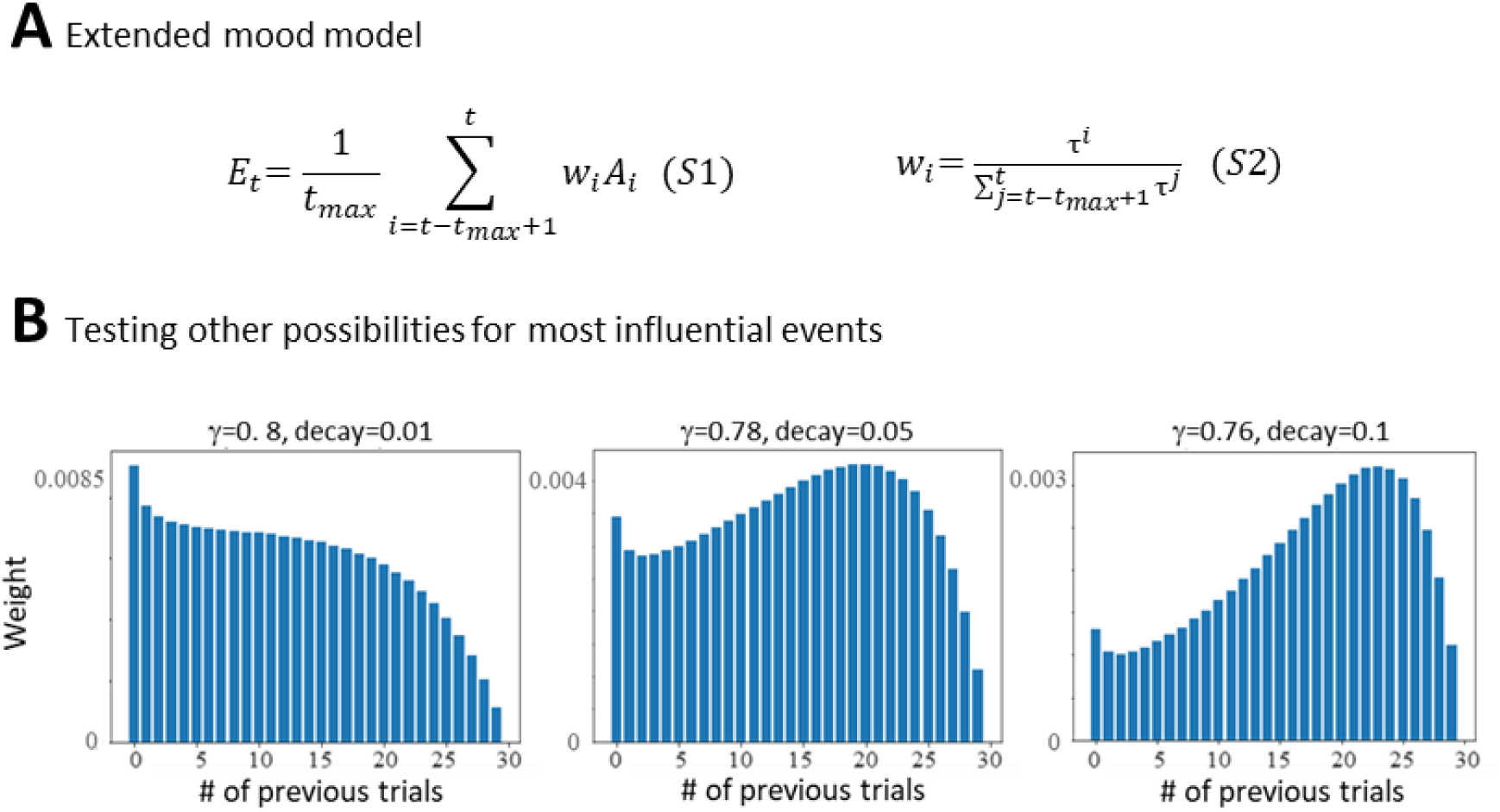
Expanding the Primacy model. (A) Eq. S1,S2 show the two additional parameters that were included in the Primacy model to test other distributions of most influential events on mood. (B) Examples for theoretical distributions that were tested with the extended model which were also outperformed by the Primacy model.

**Table S1:**
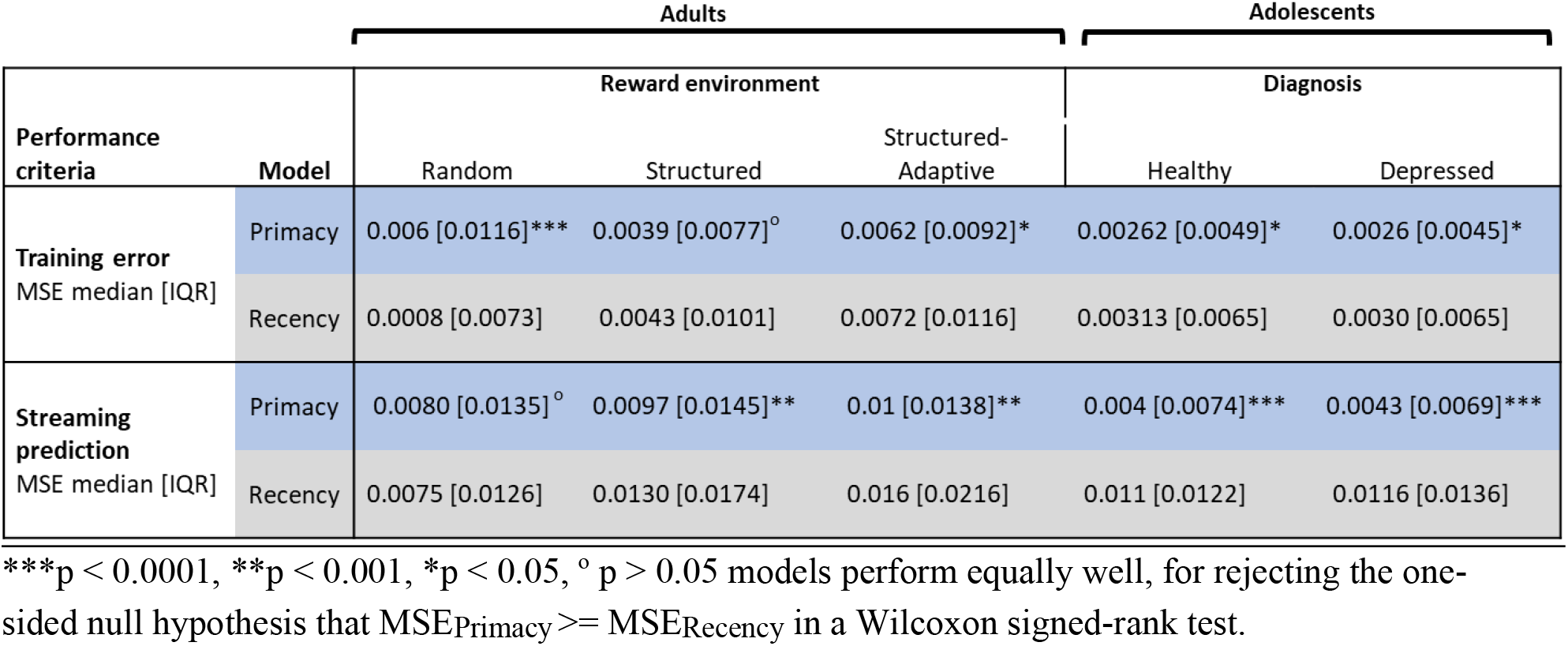
Performance of the Primacy versus Recency model, across all tasks and datasets:

**Fig. S2.**
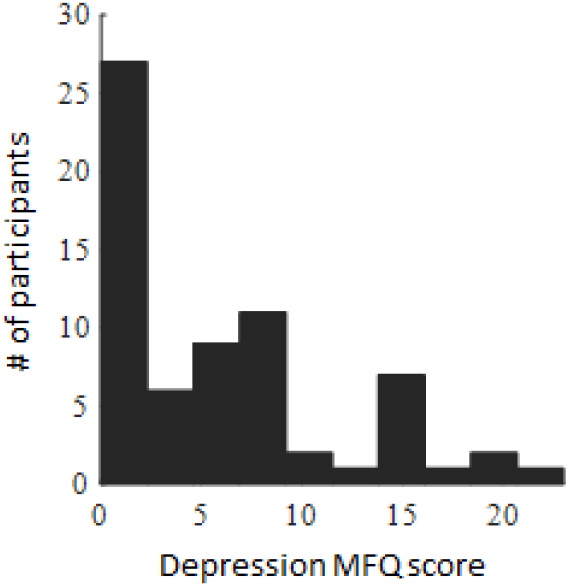
Distribution of Depression scores (MFQ) across 72 lab-based adolescent participants.

**Fig. S3.**
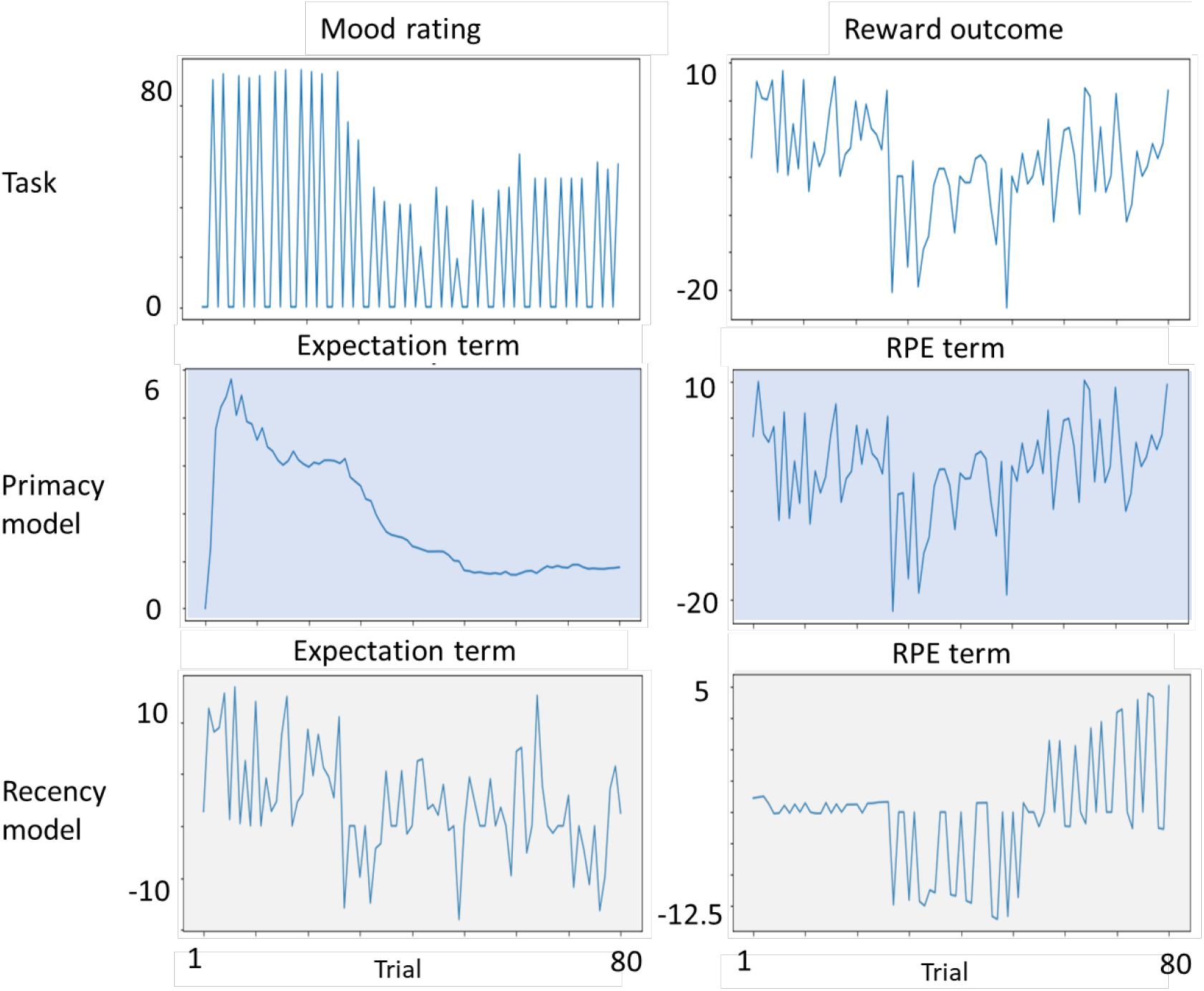
Task and model parameters of a single participant along the Structured-Adaptive task

